# Combining EEG and Eye-Tracking in Virtual Reality - Obtaining Fixation-Onset ERPs and ERSPs

**DOI:** 10.1101/2024.04.24.590885

**Authors:** Debora Nolte, Marc Vidal De Palol, Ashima Keshava, John Madrid-Carvajal, Anna L. Gert, Eva-Marie von Butler, Pelin Kömürlüoğlu, Peter König

## Abstract

Extensive research conducted in controlled laboratory settings has prompted an inquiry into how results can be generalized to real-world situations influenced by the subjects’ actions. Virtual reality lends itself ideally to investigating complex situations but requires accurate classification of eye movements, especially when combining it with time-sensitive data such as EEG. We recorded eye-tracking data in virtual reality and classified it into gazes and saccades using a velocity-based classification algorithm, and we cut the continuous data into smaller segments to deal with varying noise levels, as introduced in the REMoDNav algorithm. Furthermore, we corrected for participants’ translational movement in virtual reality. Various measures, including visual inspection, event durations, and the velocity and dispersion distributions before and after gaze onset, indicate that we can accurately classify the continuous, free-exploration data. Combining the classified eye-tracking with the EEG data, we generated fixation-onset ERPs and ERSPs, providing further evidence for the quality of the eye movement classification and timing of the onset of events. Finally, investigating the correlation between single trials and the average ERP and ERSP identified that fixation-onset ERSPs are less time-sensitive, require fewer repetitions of the same behavior, and are potentially better suited to study EEG signatures in naturalistic settings. We modified, designed, and tested an algorithm that allows the combination of EEG and eye-tracking data recorded in virtual reality.

---

Decades of research conducted in traditional laboratory settings have raised the question of how real-world contexts and conditions may impact the validity and applicability of current results (e.g., Makeig et al., 2009; Matusz et al., 2019; Shamay-Tsoory & Mendelsohn, 2019; Bohil et al., 2011). Moving out of the lab and into real-world settings provides many challenges, such as low reproducibility and limited information on the collected data (Shamay-Tsoory & Mendelsohn, 2019), that might make drawing conclusions difficult. The continuing strive to investigate human behavior beyond the classical lab setups, the challenge of real-world experiments, and a rise in new and better technology have led many researchers to utilize virtual reality (VR) to bridge this gap (Bell et al., 2020; Draschkow et al., 2022; Helbing et al., 2020, 2022; Llanes-Jurado et al., 2020). Virtual reality allows for complex experiments with a high degree of freedom for the subject while maintaining high control and replicability and providing knowledge about participants’ behavior (Bohil et al., 2011; Pan & Hamilton, 2018). As a result, virtual reality is a promising route to investigating behavior in situations of varying complexity.

Still, to fully utilize subjects’ free viewing and free exploration behavior, methods like eye movement classifications must be employed in VR experiments. While much work has been invested in classifying eye movements in two-dimensional environments, leading to well-defined, well-tested, and applied eye-tracking algorithms (Andersson et al., 2017), the literature still needs to reach this stage for three-dimensional environments like those present in VR studies. Recording eye-tracking data in the much-used Unity3D VR environment additionally provides the challenge of a low and comparatively unstable sampling rate. However, good progress has been made recently. Eye-tracking algorithms introduced for three-dimensional free-viewing data can be grouped into velocity and dispersion-based algorithms (Duchowski, 2017; Llanes-Jurado et al., 2020), similar to two-dimensional algorithms. For example, Llanes-Jurado and colleagues (2020) propose a dispersion-based algorithm suitable for three-dimensional data recorded in Unity3D. Similarly, Keshava and colleagues (2023) use a velocity-based algorithm adjusted to fit trial-based free-viewing data recorded in VR. For classifying continuous data with varying noise levels, making it difficult to detect small eye movements, Dar and colleagues (2021) offer the solution to split the recording into smaller segments based on high spikes of noisy velocity data. This velocity-based algorithm has yet to be tested for data recorded in VR. Furthermore, subjects’ eye and body movements challenge the calculation of eye movement velocities. Recording eye movements in three-dimensional VR scenes while allowing for rotational or translational movements results in eye-in-head movements or rotations of the eye in the world that do not directly correspond to movements of the optical image on the retina. Moreover, accounting and correction for subjects’ movements are essential for reliably using velocity-based eye movement classification algorithms. Employing a velocity-based algorithm that can classify continuous data allows virtual reality studies employing free viewing and exploration behavior to be implemented and compared to the existing literature.

Additional complexity is added when recording EEG in free-viewing virtual reality studies. In traditional laboratory-based setups, the timing of stimulus presentation, while the subject is fixating on a display, is under the precise control of the experimenter and can be used to align different trials across long recordings. The absence of predefined trials with repetitions of behavior makes drawing conclusions and subject comparisons difficult (Dar et al., 2021). In free-viewing experiments, this challenge can be circumvented by defining event or fixation onsets as equivalent to trial onsets (Dimigen et al., 2011), requiring eye-tracking algorithms to capture these accurately and precisely. These allow for the analysis of fixation-onset event-related potentials (fERPs) and event-related spectral perturbations (fERSPs) (Dimigen et al., 2011; Gert et al., 2022), making eye movements classification, including their precise timing, critical. Therefore, as they are better suited to determine the saccade offsets and gaze onsets (Nyström & Holmqvist, 2010), velocity-based algorithms are preferred to be combined with time-sensitive measures, such as EEG. Furthermore, the complexities of measuring eye movements while exploring a 3D virtual environment interact to define the temporal onset of fixation events precisely. To our knowledge, an eye-tracking algorithm suitable to analyze fixation-onset ERP/ERSPs in free-viewing virtual reality experiments while accurately accounting for subject movement has yet to be implemented and tested.

In the present study, we aim to develop the procedures and validate recordings of EEG and eye movements in VR to analyze fERPs and fERSPs. We created a virtual, three-dimensional environment for our participants to explore while recording eye-tracking and EEG. Eye movements were classified using a velocity-based algorithm and an adaptively determined velocity threshold (Keshava et al., 2023; Voloh et al., 2020).

Furthermore, we used a data-driven segmentation method to deal with long recordings without predefined trials (Dar et al., 2021). As our subjects could move freely within the virtual scene, we corrected their translational movements, ensuring a sound interpretation of eye movement velocities. Note that our algorithm does not differentiate between different types of eye-stabilization movements but instead summarizes them as “gazes”. The output of our data classification was tested via visual inspection and various measures, including event duration, temporal development of changes in velocities, and the comparison with hand-labeled data. We then used the gaze onsets as event and trial onset and investigated the quality of the EEG signal. Specifically, we investigated fERPs and fERSPs, the first of which, measuring evoked activity, requires a precise trial onset (Luck, 2014), and the second, while less time-sensitive, still relies on the onset of events (Cohen, 2014). Finally, we compared the EEG signature of each trial to the average fERP and fERSP signal. We observed a lower time sensitivity and a lower variance over trials for ERSPs compared to ERPs, indicating that fERSPs are potentially better suited for free viewing and exploration experiments. Overall, we can accurately determine saccade offset or gaze onset using the eye-tracking data recorded with Unity, which is suitable for analyzing fERPs and fERSPs.

## Methods

### Participants

Thirty-six subjects participated in this study. Of the 36 recorded subjects, 17 had to be rejected for various reasons: seven due to noisy EEG signals (determined by the amount of data rejected), three subjects due to failure to accurately perform the eye-tracking validation, one subject due to an inaccurate sampling rate, and four due to motion sickness. Two subjects were rejected as they failed to stay within the designated movement area, even after repeated reminders. After rejecting subjects, the final data analysis included 19 subjects (ten females, zero divers, mean age = 21.737 ± 1.408 years), predominantly students from the University of Osnabrück. All 19 subjects had normal or corrected-to-normal vision and gave written informed consent to participate in the study. Their participation was rewarded with monetary compensation or course credits. The ethics committee of the University of Osnabrück approved the study.

### Experimental Setup and Procedure

The virtual environment was built using Unity3D (www.unity.com) version 2019.4.21f1 using the built-in Universal Render Pipeline/Unlit and one central light source to illuminate the environment uniformly. The three-dimensional city center-type environment comprised 118 houses, background objects such as trees or park benches, and different pedestrians (see Fig.1; prefabs taken from Nezami et al., 2020). During the experiment, participants wore the HTC ViveProEye HMD (110° field of view, resolution 1440 x 1600 pixels per eye, refresh rate 90 Hz; https://business.vive.com/us/product/vive-pro-eye-office/) with the Eye-Tracking SDK SRanipal (v1.1.0.1; https://developer-express.vive.com/resources/vive-sense/eye-and-facial-tracking-sdk/). Movements were tracked using the HTC Vive Lighthouse 2.0 tracking system (https://www.vive.com/eu/accessory/base-station2/). Moving forward and backward within the virtual scene was made possible by using the HTC Vive controllers 2.0 (https://www.vive.com/eu/accessory/controller2018/, sensory feedback disabled). The experiment was displayed at a constant frame rate of 90 Hz, recorded on an Alienware Aurora Ryzen computer (using a Windows 10 system, 64-bit, build-version 19044; 6553 MB RAM) with an Nvidia RTX 3090 GPU (driver version 31.0.15.2698) and an AMD Ryzen 9 3900X 12-Core CPU. Behavioral data, including eye-tracking data and participant movement, were saved using Unity’s ‘FixedUpdate’ loop. EEG data were recorded using the OpenVIBE acquisition server, v2.2.0 (Renard et al., 2010; see ‘EEG data acquisition and analysis’ for more details). This was done on a separate Dell Inc. Precision 5820 Tower computer using a Windows 10 system (64-bit, Build 19044), an Nvidia RTX 2080 Ti GPU (driver version 31.0.15.1694), and an Intel Xeon W-2133 CPU. EEG and behavioral data were aligned with LabStreamingLayer (LSL; https://github.com/sccn/labstreaminglayer). During the experiment, subjects were seated on a wooden swivel chair that allowed them to turn 360° (see Fig.1A). The EEG amplifier was placed in a wooden box on the back of the chair.

**Figure 1.**
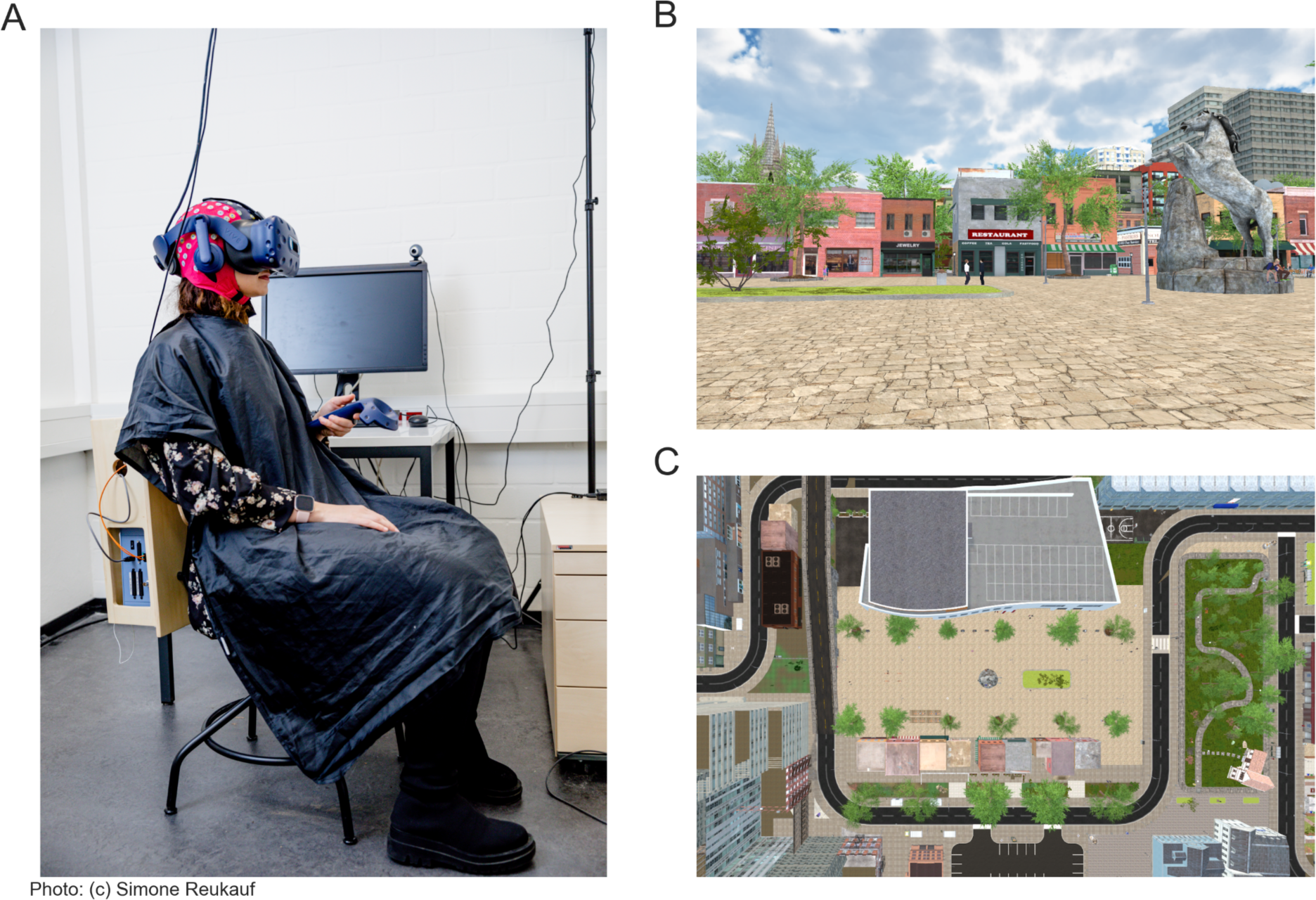
Experimental Setup and the Virtual Reality Scene. **(A)** Experimental setup with the participant wearing an EEG cap and VR glasses. The participant is seated on a swivel chair with the amplifier placed in a wooden box on the back of the chair. **(B)** A scene of the virtual reality scene. **(C)** The bird’s eye view of the central section of the city. The gray round dot in the middle is the horse statue, and the small squares on the lower half of the screen are the roofs of the houses seen in **(B)**.

The experiment was designed as a free exploration study with minimal task instructions: Subjects should imagine waiting for a friend they plan to meet. They were consequently instructed to stay close by and not leave the beige pavement (see Fig. 1B and C). This was the only area where houses and other city-center-like structures surrounded the subjects. Additionally, we gave the subjects explicit instructions to behave as close to real life as possible, with the exception of not rotating their heads up or down, as our pilot recordings revealed that this movement elicited noisy EEG data. Within the designated area, subjects could move freely. The subjects’ movement consisted of small displacements, mimicking real-life walking movements. After receiving the experimental instruction, subjects started with a five-point eye-tracking calibration and validation test, followed by a one-minute-long motion sickness test, completed in the same virtual city but in a separate street that could not be reached during the experiment. Only when passing these tests were subjects allowed to participate in the study. The experimental duration was 30 minutes, with 5-point validation and possible re-calibration break every five minutes.

### Temporal alignment checks of data streams using LabStreamingLayer

As timestamps were recorded and sent to LabStreamingLayer via Unity3D and the OpenVIBE acquisition server, v2.2.0 (Renard et al., 2010), we implemented specific latency checks to control for timing differences that might influence our analysis. These tests were completed without subjects wearing the headsets before the actual data collection of the study began. For these tests, a plane switched in Unity once every 500 ms from black to white for precisely one frame before turning back to black. These switches between black and white were pushed to LSL via the Unity ‘FixedUpdate’ loop, using a fixed frame rate of 90 Hz. A photodiode attached to the HMD, acting as the light sensor, captured the changes once they were visible and fed them directly into the EEG amplifier at a sampling rate of 1024 Hz. The input from the photodiode was measured via the OpenVIBE acquisition server (Renard et al., 2010). Six repetitions of these latency tests were performed, recording durations varying from 7.32 to 65.80 minutes (median = 20.49 minutes; IQR = 12.9-20.64 minutes). These latency tests were performed four times in the virtual environment used for recording and twice in a two-dimensional environment built explicitly for latency tests (for the analysis, see Vidal De Palol & Nolte, 2020). The recorded latencies are the differences between the measured timestamps of Unity and EEG over time and across recordings. These steps can be used to assess the quality of the temporal alignment of the different recorded streams.

A linear drift between the time stamps recorded by the OpenVibe acquisition server (Renard et al., 2010) and Unity occurred during the recordings. The absolute difference between EEG and Unity timestamps between the beginning and end of the recording (median across subjects = 53 ms; IQR = 45 ms - 87 ms) was added to the recorded timestamps to correct this drift for each participant separately. In order to account for possible inaccuracies of frame drops from Unity, the Unity time stamps were interpolated before the drift was linearly added to the data during analysis. The interpolated timestamps were excluded before defining the event and therefore the onsets used for the EEG analysis. Additionally, a shift between the beginning of the EEG and Unity timestamps was removed by subtracting the first EEG timestamp from the Unity timestamps. Importantly, to accurately align the different time streams, both eye-tracking and EEG should be recorded from the start of the experiment until the very end. We applied LSL to synchronize the different devices and data streams. Additionally, we corrected the procedural drifts by a linear fit. Specifically, we determined the change of duration at the start and end of the different streams and corrected this drift using the time axis only. Please note that this correction used no information from the recorded EEG or eye-tracking data. As a result, the two timestreams could be accurately aligned.

### Eye-tracking preprocessing and classification

Before applying the eye-tracking classification algorithm, the data was preprocessed. Invalid eye-tracking samples, which included blinks, were recorded and marked in the data. If the duration of the invalid period was longer than 20 ms, samples up to 23 ms before and after the invalid period were also rejected (Dar et al., 2021). Afterward, invalid periods smaller than 250 ms were interpolated (median amount of interpolated data across subjects = 2.823%; IQR = 1.881% - 3.914%). This rather long period was chosen in accordance with Walter et al. (2022). There, the interpolation period was justified with the analysis that subjects were unlikely to fixate on one object, move their fixation to another object, and then back again within the period of 250 ms (Walter et al., 2022). In the current paper, while we interpolated long data segments, invalid data is still rejected from further analysis.

The participant positions and hit points (the intersection of the gaze direction vectors with objects in the world) needed for the eye-movement classification, specifically the translational movement correction, were filtered using a 5-point median filter. We selected this filtering method as it deals well with single outlying samples without low-passing, smoothing saccadic onset and offset. This preprocessed eye-tracking data was then used for eye movement classification.

The employed eye movement classification algorithm differentiates between fixations and saccades based on the eye angular velocity, which is influenced by translational movement within the virtual environment. Figure 2A displays two different hit points that might be viewed in sequence, one on the ground (hit point 1) and one on a building (hit point 2), as well as two different subject positions (subject positions 1 and 2) between which a translational movement occurred. Note that the distance between subject positions 1 and 2 has been exaggerated relative to the distance of the viewed objects for didactic purposes. This graph demonstrates the two potential issues that arise when not correcting for translational movement. First, one can consider that a translational movement occurs while the subject focuses on the same point in the world. When viewing hit point 2, the green (diagonal) vector indicates the allocentric eye direction at the first point in time. The red (straight up) vector indicates the eye direction at the second point in time. If not correcting for translational movement, we infer a non-zero angular velocity, i.e., the angle between the green and red vectors, even though the object viewed and the corresponding hit point stays the same. Alternatively, the subject could translate their body without changing the allocentric eye direction. This results in two different hit points but zero angular velocity of the eye. In our figure, this is represented by the parallel blue and red allocentric eye directions. If we do not correct the translation of the subject, our algorithm will not recognize a shift in hit points even though one occurred. As a result, it is essential to correct translational movement in a virtual environment that includes translational movements of the subjects.

**Figure 2.**
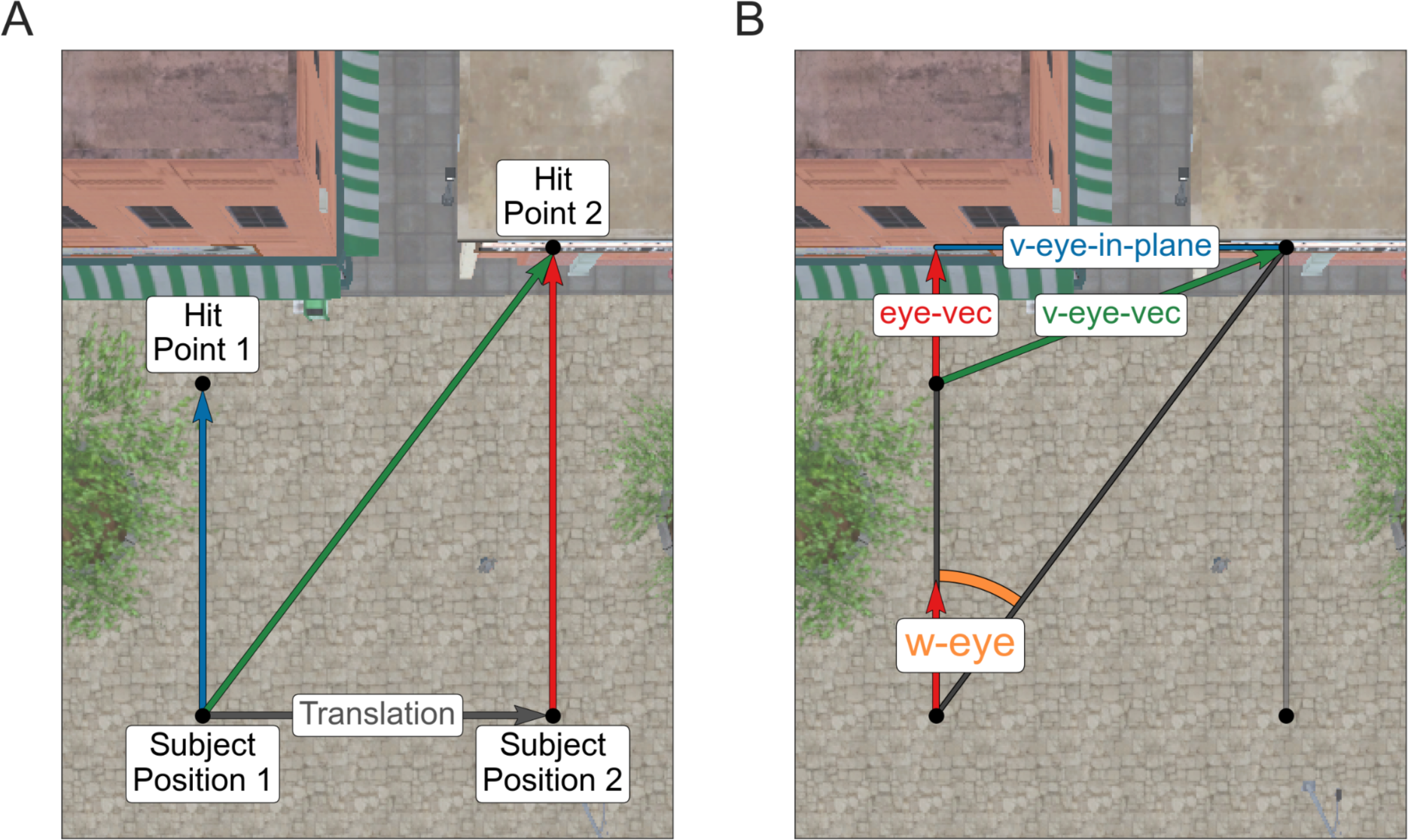
Schematic of the Translational Movement Correction. Seen here is a schematic overview of our correction for translational movements. The plot **(A)** motivates using a translational movement correction within virtual reality. We show two consecutive subject positions (subject positions 1 and 2) and a vector indicating the translation between the two and two different hit points (hit points 1 and 2) on top of a screenshot of our virtual city. Note that the distance between subject positions 1 and 2 has been exaggerated relative to the distance to the viewed objects for didactic purposes. Three possible allocentric eye directions are displayed in blue (the left of the two parallel vectors), green (diagonal), and red (the right of the two parallel vectors). Without movement corrections, an angular velocity between each combination of these vectors could occur. However, when considering the green and red vectors, where the eye direction changes but the hit position does not, or the blue and red vectors, where the hit positions change but the eye directions do not, a correction for the translation is needed to determine the differences between fixations and saccades accurately. In **(B),** we show a visual representation of the algorithm in the case of a translation but no change in the allocentric eye direction. The dark gray lines correspond to the two eye directions if the subject had not moved. While corresponding to the second viewing direction, the light gray line did not enter the equation. The two red vectors correspond to the eye direction (eye-vec), the green vector, v-eye-vec, to the shift of the hit points without taking the subject position into consideration, and the blue line v-eye-in-plane is the projection of v-eye-vec onto a plane orthogonal to the viewing direction. Finally, the angle w-eye (drawn in purple) represents the calculated angular velocity irrespective of translational movements.

We, therefore, introduced a movement correction before calculating the eye angular velocity (Keshava et al., 2023) used to define gazes and saccades. The key to our approach is first to compute the hit point’s movement in allocentric coordinates and then translate it to the required change of eye direction in allocentric coordinates. Figure 2B demonstrates this movement correction for both issues. The first step was to calculate the shift of the hit points without considering the position it was seen from, which is represented as the green vector v-eye-vec in Figure 2B. If the distal hit point did not move (first problem), this shift is zero and will stay so in the following operations. Next, this vector (v-eye-vec) was projected on a plane orthogonal to the viewing direction (eye-vec, the red vector in Fig. 2B), which we called v-eye-in-plane and represented by the blue line:

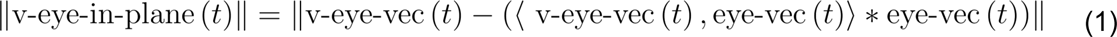

where t indicates a time point, 〈·,·〉 the scalar product, and ||…|| the Euclidean norm. Using v-eye-in-plane, we determined the changes of the eye direction in allocentric coordinates, or more specifically, the angular velocity w-eye, by calculating the arctan between v-eye-in-plane and the current viewing direction, eye-vec, and dividing this by the difference in time:

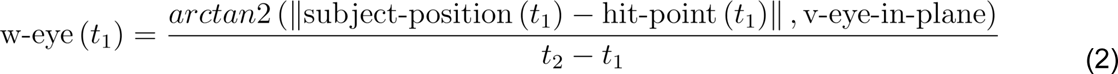

with *t*_1_ and *t*_2_ representing two consecutive time points. Finally, this angle was transformed from radians into degrees. As a result, we received the change of eye direction the subject would have made from the first to the second hit point without a translational movement. Specifically, whether the subject was standing in place while executing a change in the viewing direction or translating a considerable amount while maintaining an allocentric eye direction did not enter the calculation, thereby giving invariant results under different degrees of translation. It is important to note that the original normalized direction vectors obtained from Unity did not have a length of one. We assume this was due to an internal rounding error. In order to avoid systematic errors, it is recommended to recompute and normalize these offline.

Angular velocities above 1000 deg/sec (Dar et al., 2021) were corrected to 1000 deg/sec to exclude biologically impossible eye movements. The velocities were then filtered again using a Savitzky–Golay filter (length of 3s, polynomial order of 2).

The calculated angular velocities (Keshava et al., 2023) were classified into gaze events (smooth pursuit and fixations) and saccades based on an adaptively calculated threshold (Voloh et al., 2020). This experiment did not have predefined trials, so applying the adaptive threshold method to the entire dataset would only detect very big saccades. As a solution, data were segmented using the method introduced by Dar and colleagues (2021): data segments of the eye movement data were calculated by applying the MAD saccade algorithm to the entire eye-tracking recording to determine a global threshold. Large eye movements, typically saccades, exceeding this threshold were ranked according to the sum of velocities within this threshold and used as event boundaries until an average frequency of 2Hz across the entire recording was reached (Dar et al., 2021). Overall, across subjects, the method resulted in a median segment duration of 256 ms (mean = 251 ms; min = 167 ms; max = 299 ms). This data segmentation method is called the ‘data-driven method’ in the following text.

A preliminary analysis found that a data-driven method occasionally produced very short intervals. This depended on the filtering details and was most likely caused by our relatively low sampling rate. Therefore, for the first part of the paper, we additionally contrasted the data-driven method with the method of cutting the data into 10-second intervals. A separate threshold was calculated for each of these segments. This solution is called the ‘10-second method’ in the following. By comparing the two methods, we want to investigate the reliability of a robust data segmentation with a fixed time scale versus an adaptive algorithm.

Finally, we corrected for outliers or inconsistencies within the data. If the direction vector hit more than one object during the same gaze interval, the object with the most hits was considered the current focus. Events with biologically implausible durations (saccades<20ms, gazes<40ms) were merged with the previous event. The slightly longer minimum saccade duration of 20 ms compared to the literature (Dar, Asim H. et al., 2021) was due to the relatively low sampling rate of 90 Hz. In line with Keshava et a. (2023), too-long gazes or saccades were identified and rejected if they deviated more than 3.5 from the median absolute deviation across all subjects. Finally, invalid eye-tracking periods were rejected. The classification for one subject is shown in Figure 3. In the paper, we use this same subject to demonstrate the results for a single subject. The subject was not selected for specific reasons other than being the first subject recorded.

**Figure 3.**
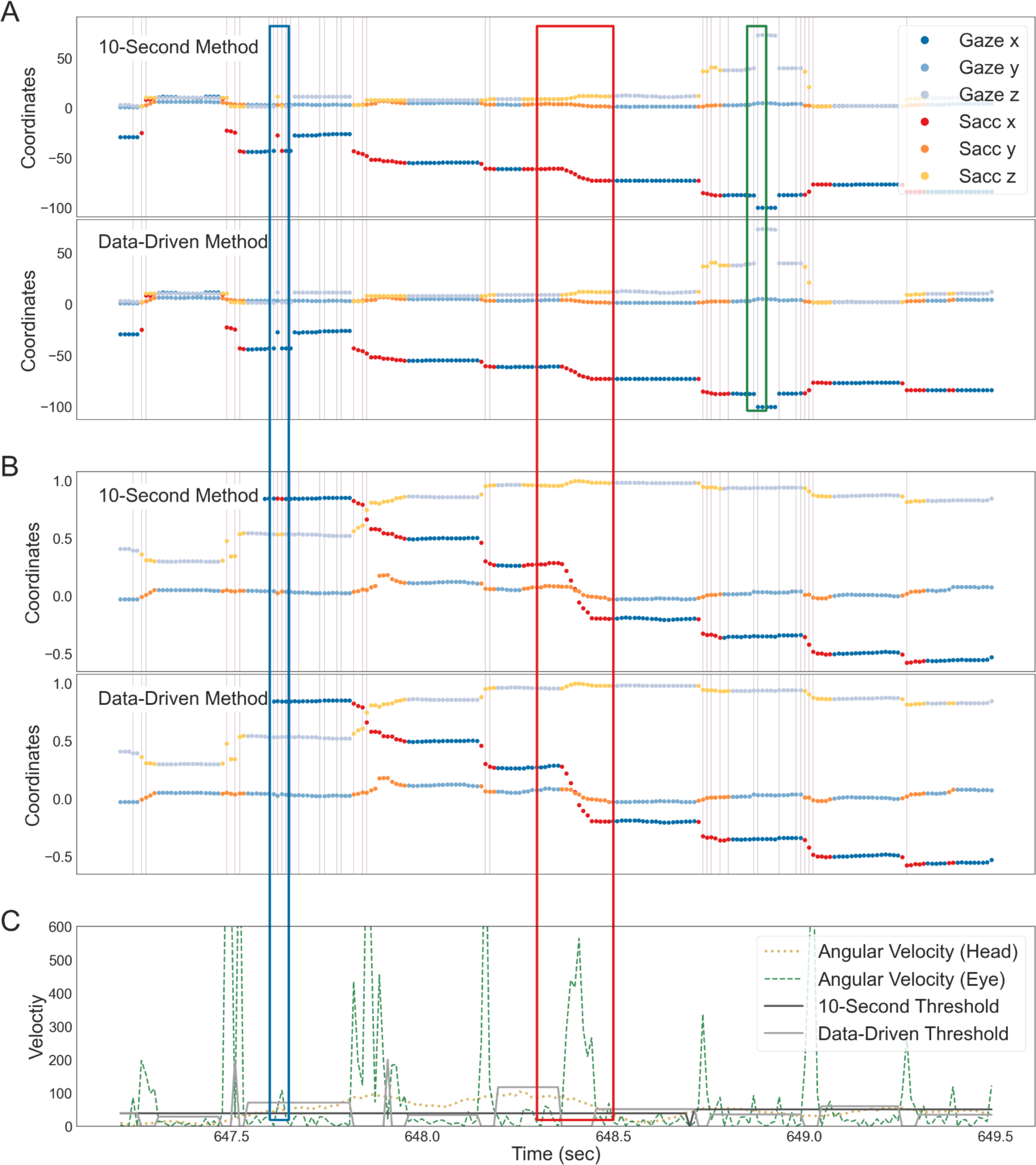
Results of the Eye Tracking Classifications. These data were selected to show various properties and problems of the algorithm. **(A)** and **(B)**: The output of both data segmentation methods is shown. The x-,y-, and z-coordinates for hit points **(A)** and direction vectors **(B)** are plotted against time (sec). Each coordinate is divided into gaze (in blue) and saccade (red-yellow colors) events. Vertical lines indicate that the eye direction hit a new collider. In **(C)**, the eye angular velocity (dashed green line), as well as the head angular velocity (light green dotted line), are plotted over the same period. The classification threshold for the 10-second method is shown in dark and the data-driven method in light grey. The colored outlined rectangles indicate events of interest (see the result section ‘Classification fit’ for more details).

To assess the accuracy of this eye-tracking algorithm, we compared the output of the classification algorithm to the hand-labeled data of one subject. We manually classified gazes and saccades for the first subject using the unfiltered gaze direction vectors. Additionally, we classified invalid data using the information given by the eye-tracker. Analogous to the algorithm-defined events, we corrected for outliers in the manually classified data if their duration deviated more than 3.5 from the median absolute deviation. The results of the hand-labeled data were compared to the algorithm performance on a sample-by-sample and event-onset basis.

### Data distribution

The data distribution was assessed by analyzing the number and percentages of gazes, saccades, and outliers. The number of gazes and saccades was calculated separately for both data segmentation algorithms. As the counts across subjects were not normally distributed, we report the median and interquartile range (IQR = Q1 - Q3). Differences in the number of gazes and saccades between the segmentation algorithms were quantified using a Kolmogorov-Smirnov (KS) test. Similarly, percentages of events were calculated for both classification algorithms, excluding breaks for validation and calibration. Outliers were divided into invalid samples, composed of blinks and the inability to accurately detect the eyes during the recording, and outliers of event durations that were too long, reported separately for gazes and saccades. Differences between the data segmentation algorithms for percentages were assessed using a KS test and a post hoc Bonferroni correction. Besides the visual inspection, these are the only analyses used to compare the two data-segmentation algorithms.

### EEG data acquisition and analysis

We recorded EEG data with a 64-channel Ag/AgCl-electrode system, placed according to the international 10/20 system using a Waveguard cap (ANT, Netherlands) and a Refa8 (TMSi, Netherlands) amplifier. Impedances were kept below 10 kΩ. Data was recorded at 1024 Hz using an average reference and a ground electrode placed under the left collarbone. The data was preprocessed using EEGLAB (Delorme & Makeig, 2004). First, the data was re-referenced to Cz for preprocessing, high pass filtered to 0.5 Hz (EEGLAB plugin firfilt, pop_eegfiltnew, using a hamming window; Widmann et al., 2015), and downsampled to 512 Hz. As a note, we did not apply a baseline correction, as there is no obvious choice of baseline in our dataset consisting of a sequence of gazes and saccades occurring in alternation. As a remedy, we applied a relatively high band limit of 0.5Hz. Gaze onsets defined using the eye-tracking algorithm and data-driven segmentation method were inserted into the data. Noisy channels (mean channels rejected = 13.89 ± 4.74) were visually inspected and channels with comparatively high power in high frequencies were rejected from further analysis. Additionally, if individual channels were noisy over longer segments and the rejection of those segments would have resulted in a rejection of the entire subject, these noisy channels were also rejected. Similarly, data segments containing muscle activity were visually inspected and rejected (median number of trials rejected = 1251, IQR = 863.5 - 1447.5). Subjects with more than 40% of rejected data were excluded from further analysis. Eye movements were manually rejected using independent component analysis (ICA, amica12, Palmer et al., 2012) performed on epoched and high-pass filtered data at 2 Hz (Dimigen, 2020; mean ICs rejected = 14.78 ± 3.7), guided by the output of ICLabel (Pion-Tonachini et al., 2019). After applying the ICA weights on the continuous data, the data was re-referenced to average reference, and noisy channels were interpolated (spherical interpolation). For EEG data segmentation, we considered fixation-onset defined events within the spirit of classical stimulus-driven setups. In the following, when speaking of one trial (terminology adapted from standard ERP experiments), we refer to a segment of the EEG signal lasting from -200 up to 500 ms around a gaze onset. Therefore, the only results of the eye-tracking classification algorithms relevant for the EEG data analysis, such as ERPs or ERSPs, are the exact timing of the gaze onsets.

For the time-frequency analysis, the preprocessed data was epoched from -875 ms to 1175 ms, leaving a buffer zone of three cycles of 8 Hz to avoid contamination of the results with edge effects later on (Cohen, 2014). The time-frequency decomposition was calculated with the FieldTrip (Oostenveld et al., 2011) function *ft_freqanalysis* using a Morlet wavelet. Since the trade-off between the time and frequency resolution can be controlled with the width of the wavelet (Cohen, 2014; Gross, 2014), the wavelets consistently had three cycles, favoring a better time resolution. The decision to favor a time over a frequency resolution was made to evaluate the effects of the precise timing of fixation-onset defined events on the measured neuronal activity. The time-frequency analysis was conducted for single subjects and all subjects. For the time-frequency analysis, we compare the power in different frequency bands before and after the onset of an eye movement event. Considering the average saccade durations observed for the subjects, which were predominantly shorter than 200 ms, a baseline period ranging from -500 ms to -200 ms was chosen to reduce any potential bias caused by saccadic eye movements (Cohen, 2014) when calculating the time-frequency decomposition. While the current baseline period was chosen to exclude the specific saccadic period leading up to the gaze event, we acknowledge that the interval of 300ms will typically include other fixations and eye movements, like saccades.

## Results

### LabStreamingLayer and data stream alignments

Different tests revealed that using LSL to align timestreams from different recording computers or software is possible for our setup. We recorded a mean latency between sending and displaying a visual stimulus of 82 ms (std = 0.6 ms). Additionally, we recorded an average temporal distance between individual frames of 0.98 ms for the EEG time stream (equivalent to 1024 Hz; std = 0.000 ms) and a temporal distance of 11 ms for the Unity time streams (equivalent to 90 Hz; std = 0.9 ms). Taken together, these results confirmed constant latencies when using LSL and only a low number of frame drops after enforcing 90 FPS.

### Eye-tracking classification

We must rely on secondary measures to validate the eye-tracking algorithm, as we do not have ground truth in three-dimensional recordings. Therefore, as a first step, we visually inspected the classification results (Figure 3). Please note that the subjective impression of the classification of hit points (Fig. 3A) and eye direction vectors (Fig. 3B) as gazes or saccades were rather similar for both algorithms. Only at closer inspection were small differences visible. Both segmentation algorithms showed that changes in the hit position often, but not always (middle red outlined rectangle, Fig. 3A-C), correlated with a change in collider (displayed using purple vertical lines), meaning that eye movements were made between objects. Additionally, a change in the hit position was usually aligned with saccades, indicated by a change in the color of hit points and gaze direction. In a few cases, a change of the hit position was not accompanied by a saccade, as seen at around 648.8 s (green small outlined rectangle). These were likely due to inaccurate measurements or a too-low sampling rate of the eye-tracker to capture very small eye movements. Figure 3 also shows events corrected for being too short, e.g., at around 647.6 s (blue outlined rectangle) for the data-driven but not a 10-second method (Fig. 3A&B). Investigating the differences between the two segmentation algorithms revealed an additional deviation: while both segmentation methods classified events simultaneously, some of the saccades for the 10-second method are notably longer than the ones for the data-driven method, e.g., at 648.4 s (red outlined rectangle). This was a result of a comparatively lower classification threshold in this interval. Finally, comparing the angular head velocity and the angular eye velocity (Fig. 3C), some large saccades aligned with head movements. This means that both segmentation algorithms matched the underlying data well and classified the eye-tracking output similar to what one would expect by manual classification, with the data-driven method fitting slightly better than the 10-second interval.

### Data distribution

We investigated the distribution of gazes, saccades, and outliers to explore potentially unusual distributions and differences between the two segmentation algorithms. Counting the different events after outlier rejection, the data-driven method resulted in a median of 4917 total gazes (IQR = 4384.5 - 5105.5; median number of saccades = 4704, IQR = 4351.5 - 4863.0) and the 10-second method in 4780 total gazes (IQR = 4546.0 - 4863.0; median number of saccades = 4687, IQR = 4471.5 - 5073.5). A Kolmogorov-Smirnov test revealed no significant differences in the number of events for either data segmentation method (gazes: p = 0.808; saccades: p = 0.808). Next, we analyzed the percentages of the total time for the different categories (Fig. 5). The recorded data contained a median of 3.44% (IQR = 2.73 - 4.25) of invalid samples, meaning the eyes could not be accurately detected during these periods. After applying eye movement classification, investigating the distribution of events for both data segmentation methods revealed similar tendencies. For the data-driven method, a median 55.96% of the data were gazes (IQR = 53.97 - 57.12), 21.0% were saccades (IQR = 19.94 - 22.8), and 21.68% were outliers, comprising invalid samples, gazes that were too long (median = 10.36%, IQR = 8.25 - 11.85), and saccades that were too long (median = 8.35%, IQR = 7.57 - 9.81). Similarly, the 10-second method resulted in a median of 53.91% gazes (IQR = 51.67 - 55.28), 21.24% saccades (IQR = 18.86 - 22.04), and 24.85% outliers (invalid samples; long gazes = 12.63%, IQR = 10.23 - 13.74; long saccades = 9.32%, IQR = 8.74 - 9.98). Performing a KS test and a post hoc Bonferroni correction revealed no significant difference between these data segmentation algorithms. Overall, both algorithms resulted in nearly identical distributions of events. In the following analysis, we additionally tested both segmentation methods but did not observe any significant difference in the distribution of the various derived measures. As a result, the remaining analyses were reported using only the data-driven method.

### Eye and head movements

To assess the quality of the recorded data, we investigated eye and head movements after classification. Most global eye movements encompassed the horizontal x-z-plane with fewer gazes in the vertical direction directed to the very top or bottom of the environment (Fig. 5A). A similar but more extreme pattern could be seen for head movements with most head movements falling within the horizontal x-z-plane (Fig. 5B). Considering that the global eye-direction vectors combined head and local eye movements, these results indicated that head movements mediated the preference for the horizontal direction. This became apparent when investigating the normalized eye-in-head direction vectors. Figure 5C displays the eye-in-head direction vectors for the horizontal and vertical directions. The distribution of these eye-in-head direction vectors did not favor the horizontal over the vertical direction, with an average absolute maximum horizontal value of 0.59 ± 0.04 and a vertical value of 0.59 ± 0.05. Additionally, most of the eye-in-head direction vectors fell within the forward-facing direction. Finally, saccade vectors, normalized to saccade onset (Fig. 5D), were calculated by subtracting the coordinates at the beginning from the coordinates at the end of the saccade. They closely mirrored what could be observed for the eye-in-head direction vectors, spanning the entire space, with slightly longer saccades in the horizontal than vertical direction. Examining the eye movements revealed that the recorded data followed anticipated patterns and was suitable for further analysis.

Next, we investigated the saccadic main sequence to verify the validity of the classified eye movements. The accuracy of saccadic eye movements can be supported by demonstrating a correlation between saccadic amplitude and peak velocity (Bahill et al., 1975; Dar et al., 2021). We calculated saccade amplitudes corresponding to the angular velocities by correcting for subjects’ translational movement (see Figure 2 for more detail). Differently from velocities, the saccade amplitude was calculated between the centroids of two consecutive fixations. Furthermore, we did not divide the amplitude by time. As seen in Figure 5E, the peak velocity is positively correlated with saccade amplitude (*r*(79420) = .34, *p* = .0). These results are in line with previously reported effects (Bahill, 1975; Dar) and support the validity of the classified eye-tracking data.

Besides the direction vectors and the saccadic main sequence, the position towards and distance to the hit objects could be evaluated to verify the quality of the recorded data. The spatial distribution of these gazes can be seen in Figure 4G, shown overlaid on top of the central section of the virtual city. The locations of the hit objects captured the layout of the Unity scene. The colliders’ boundary lines, representing the outlines of objects used to define the object boundaries in Unity, of the different objects in the scene are visible due to many gazes falling on them, compared to the ground or sky. Generally, gazes were distributed across the entire scene, with most gazes landing within the area subjects were walking through during the experiment. Figure 4F shows the distribution of distances kept to the hit object. While we observed differences between subjects, calculating the median (mdn) and interquartile range (IQR = Q1 - Q3) for gazes and saccades across subjects’ medians revealed no differences between the distances kept to hit objects for gazes and saccades (gazes: mdn = 16.68 Unity units, IQR = 14.39 - 19.29; saccades: mdn = 16.48 Unity units, IQR = 14.37 - 19.09). A Kolmogorov-Smirnov (KS) test confirmed these results (p-value = 1.000). These results indicated that subjects maintained a certain distance from the objects they gazed at while avoiding looking at objects that were too far away.

**Figure 4.**
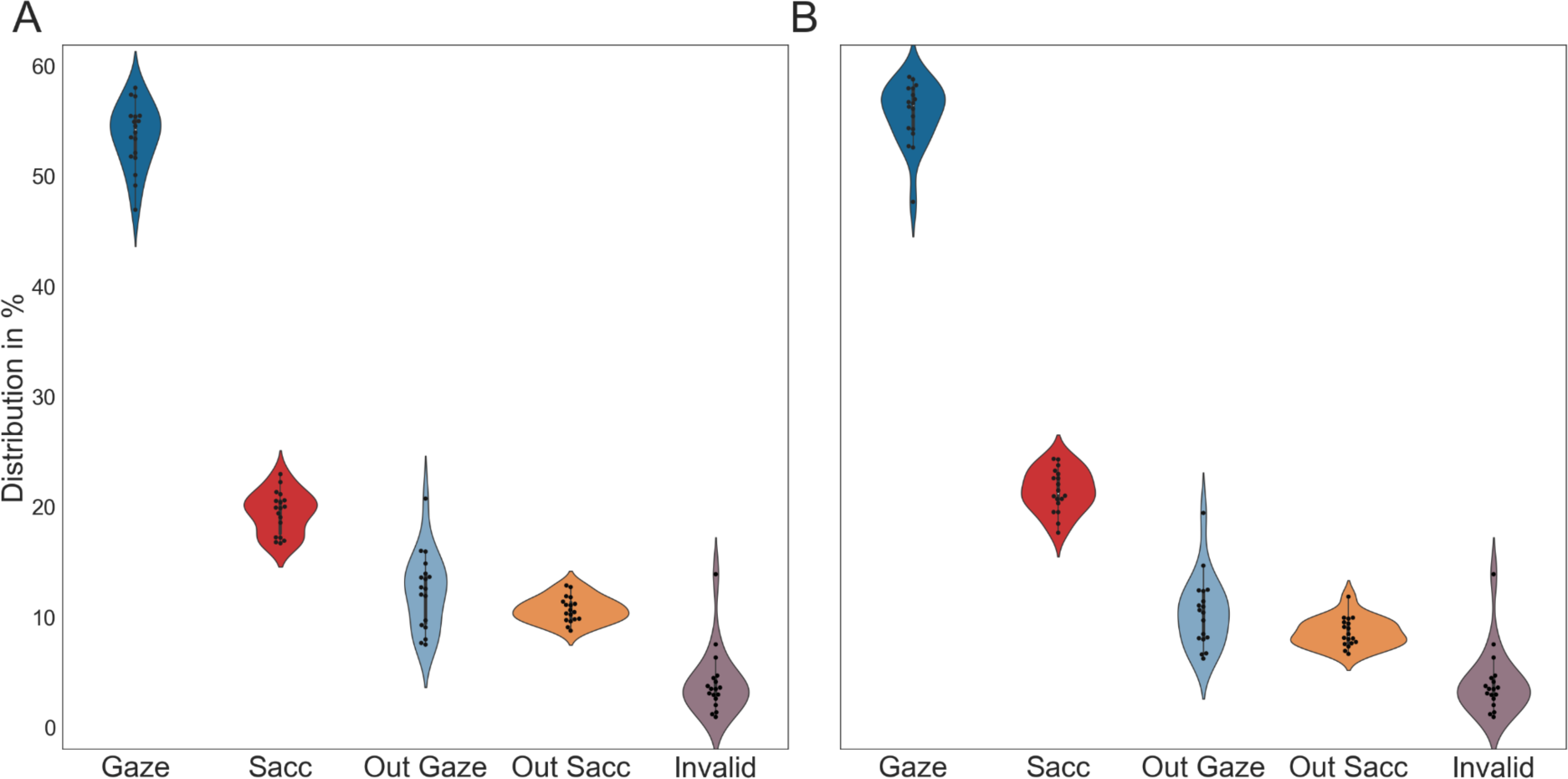
Distribution of Gazes and Saccades. The plot shows the results for both data segmentation methods, in **(A)** the 10-second method and **(B)** the data-driven method. Each data point represents the percentage of one participant. The percentages of each subject’s gazes, saccades, and outliers are calculated. ‘out gaze’ stands for gaze events excluded from the data for being too long, and ‘out sacc’ represents the percentage of saccades excluded for being too long. Invalid samples are all samples that were not accurately collected.

**Figure 5.**
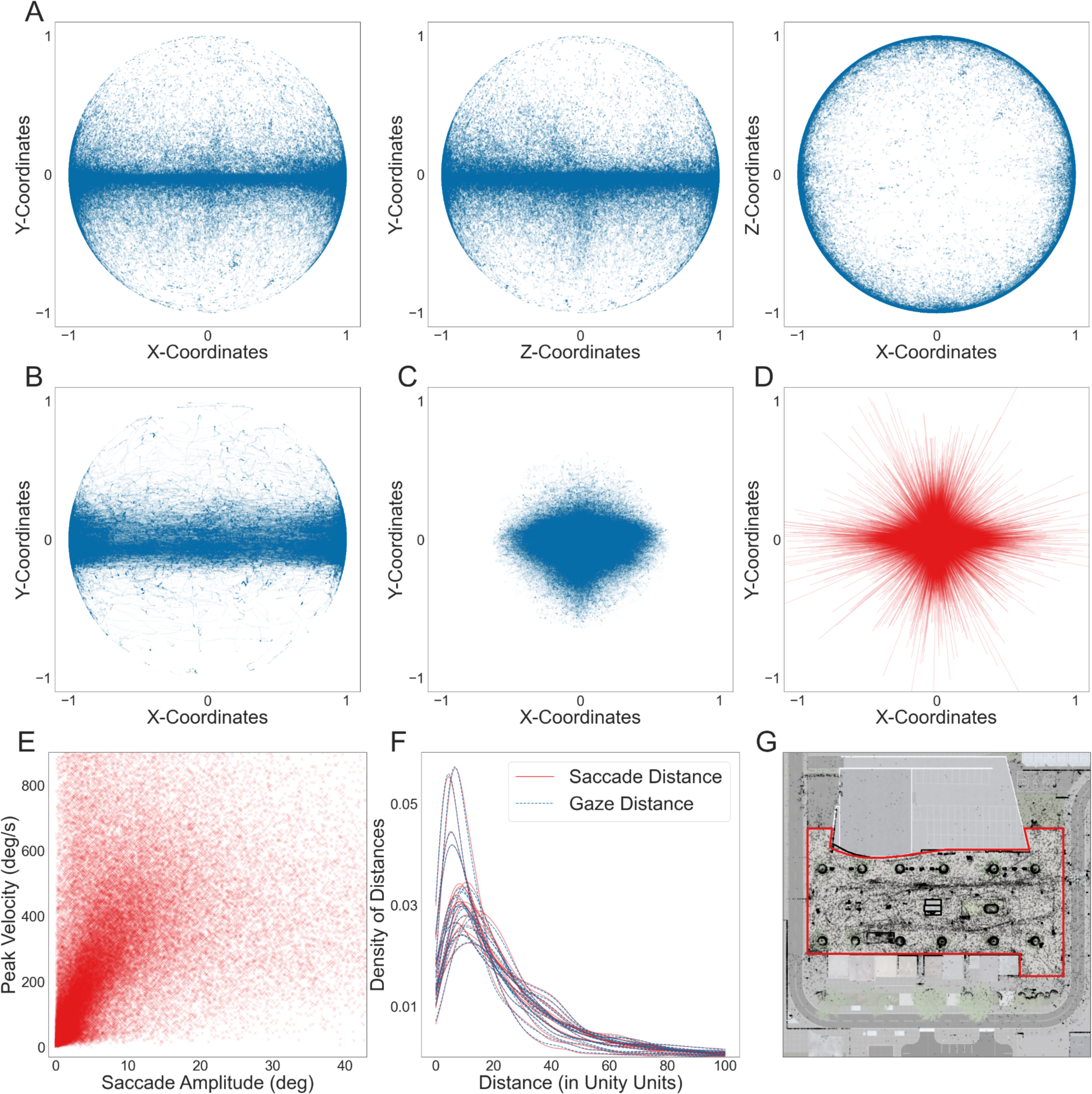
Eye and head movements. The data displayed uses the data-driven segmentation method. **(A)** Corrected and normalized direction vectors are shown for all gazes and all subjects. Each subplot depicts two of the three coordinates of the direction vector. **(B)** All participants’ normalized head direction vectors for the horizontal and vertical coordinates can be seen. **(C)** Normalized eye-in-head direction vector’s horizontal and vertical coordinates. **(D)** Horizontal and vertical coordinates of the saccade vectors aligned to saccade onset. **(E)** The peak velocity of each saccade is plotted against the saccade amplitude of the corresponding saccade. **(F)** The average distance for all subjects’ saccades (red) and gazes (blue). The density curve is plotted to start at zero, excluding impossible distances and allowing easier data comprehension. **(G)** All gazes’ average x- and z-positions are displayed for all subjects. This plot only shows the Unity scene’s central section (similar to Fig. 1C), so gazes on objects further away are not displayed. The walkable area of this plot is marked in red.

### Event durations

We investigated the average duration of saccades and gazes to validate the eye-tracking classification further. While there were minor variations among participants, the overall difference in duration between gazes and saccades remained consistent across all subjects (Fig. 6). For each subject, the duration of gazes was longer than that of saccades, and the distribution of gaze duration was broader. Across subjects, the median duration of saccades calculated by the data-driven method was 0.067 s (IQR = 0.067 - 0.068 s). Gazes had a duration of 0.178 s (IQR = 0.177 - 0.194 s). Applying a KS test revealed significant differences in the event durations (p-value = 0.0000). Additionally, means showed a similar difference between gazes (mean = 0.183 s ± 0.015) and saccades (mean = 0.07 ± 0.006). These results align with the anticipated patterns of eye movements, revealing shorter saccades and longer gazes.

**Figure 6.**
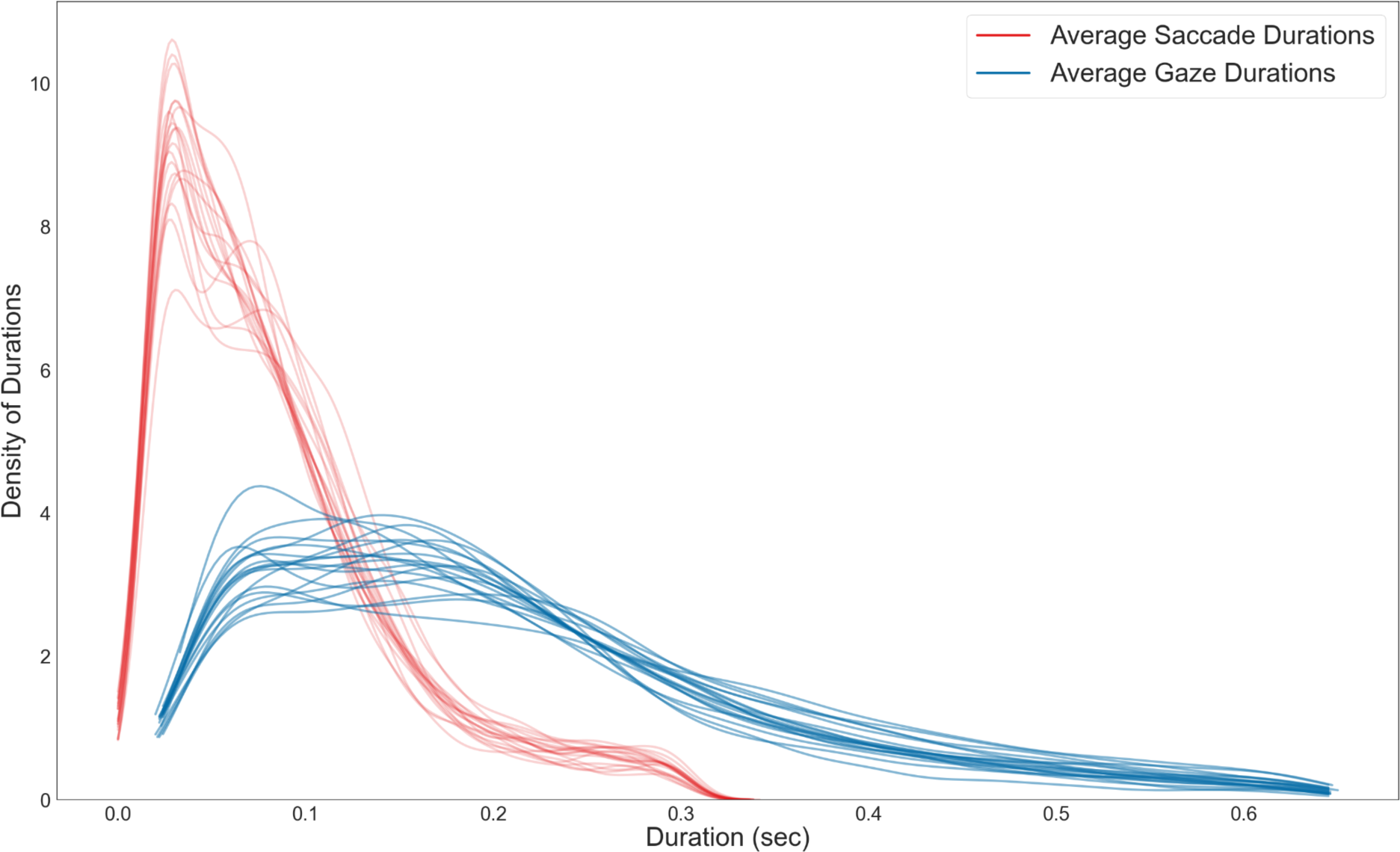
The Average. Duration **of Gazes and Saccades** The plot shows the density distributions of gaze (blue) and saccade (red) durations for all subjects individually.

### Velocity distribution in relation to gaze onset

Another way to evaluate the classification output’s quality is the velocity distribution across multiple events. The angular eye velocities before gaze onset (Fig. 7A; indicated by the red lines) were generally higher than after. Additionally, the further away from gaze onset a sample lies, the less uniform the data looked. Different peaks of high velocity occurred at different time points due to different gaze intervals potentially overlapping. Overall, for each subject, the algorithm produced high-velocity samples before gaze onset and low-velocity samples after gaze onset, as seen in Figure 7B. Velocities at about 110 ms before gaze onset slowly increased, terminating in the highest median of median velocities at about 11 ms before gaze onset. After gaze onset, the median velocities remained relatively low and stable throughout the rest of the interval, indicating less eye movement. Similarly, the distribution of peak velocities was the highest close to gaze onset, specifically at -22 to -11 ms before gaze onset (Fig. 7C). The distribution of velocities before and after gaze onset highlighted that we could seemingly define the gaze onsets as expected.

**Figure 7.**
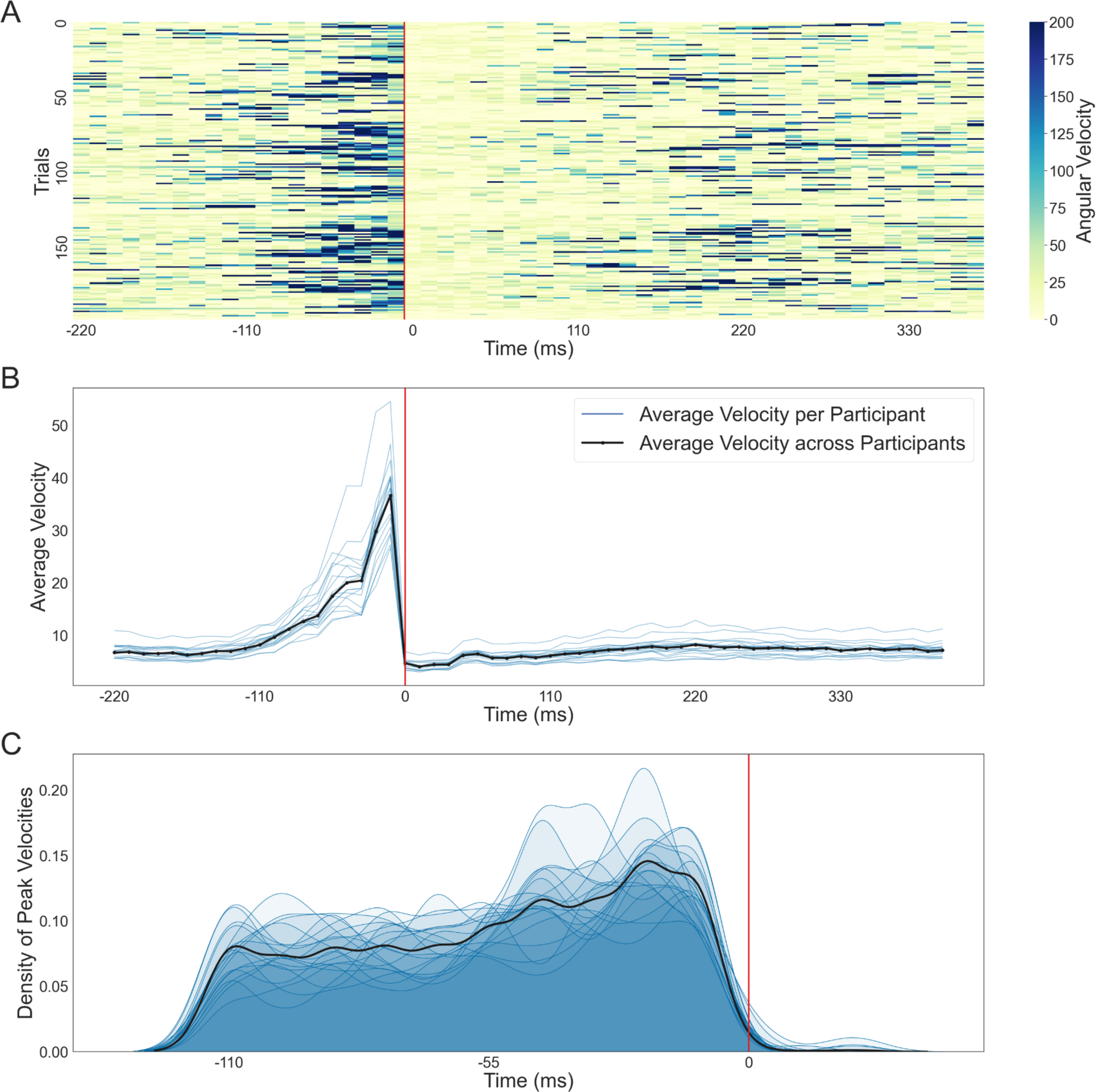
Eye Angular Velocity Distribution. **(A)** The eye’s angular velocities of 200 consecutive time periods are aligned to gaze onset (red line). Each row is centered around one gaze onset, about 220 ms before and 385 ms after gaze onset. Please note that all three subplots - **(A)**, **(B)**, and **(C)** - were created using individual frames. As a result, the time points given on the x-axis are comprised of small variations of the timing of individual values due to the slightly varying frame rate. The colors represent the different velocities, with darker blues corresponding to higher velocities. All velocities exceeding 200 degrees per second are corrected to that value for easier visibility. Depending on the length of one gaze and saccade, the end of a row can include the beginning of the next gaze. **(B)** The median velocity of each subject (blue) and the median of medians across subjects (black) plotted over time (-220 to 418 ms) are aligned to gaze onset (red line). **(C)** The density distributions of peak velocities for each subject (blue) and the distribution across all subjects (black) are shown from 110 ms before gaze onset to 22 ms after gaze onset.

### Dispersion distribution in relation to gaze onset

Besides investigating the temporal development of saccades and gazes, we studied the average dispersions for all subjects in relation to gaze onset. The change of dispersions was calculated as the Euclidean distance between two consecutive hit points. The dispersion distribution was similar to the velocity distribution. As seen in Figure 8, at roughly -110 ms before gaze onset, we saw a slow increase in the dispersion density distributions for all subjects. At about -44 ms before gaze onset, the change in the hit position reached a local peak. The locations of the hit points changed the most between the second to last and the last sample before gaze onset. After gaze onset, the density distributions remained relatively low and stable. Changes in the hit position shortly before but not after gaze onset supported a clean determination of fixation onsets.

**Figure 8.**
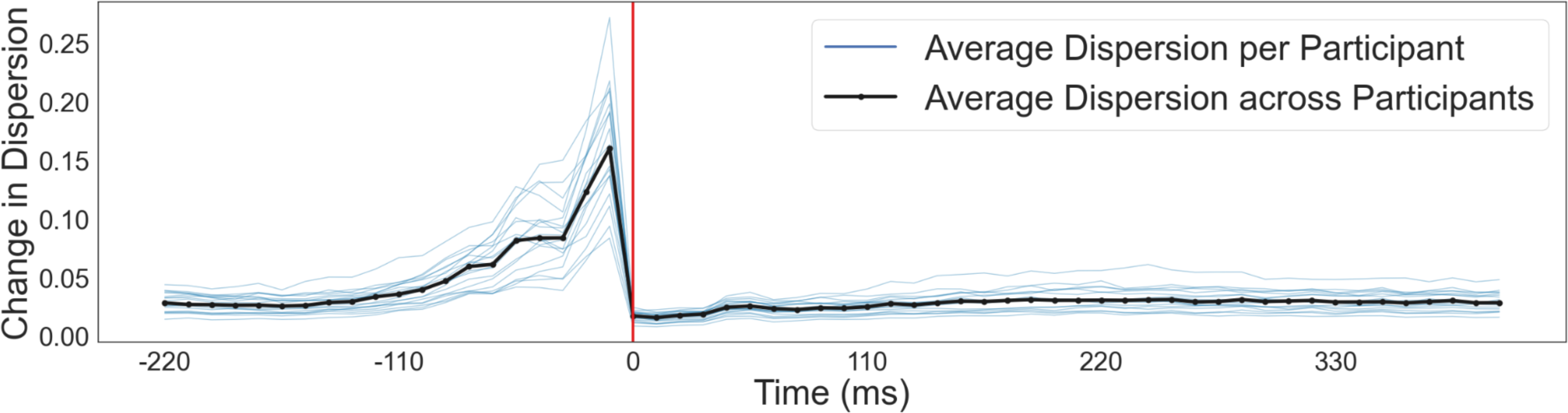
Change in the Dispersion Distribution. Similar to the velocity plots, the change in dispersion is aligned to gaze onset. The distribution is shown for roughly -220 ms before gaze onset to 418 ms after. The plot was created using frames, and as a result, the time points given on the x-axis are comprised of small variations in the timing of individual frames. The median dispersion for each subject is shown in blue, and the median across subjects is black.

## EEG Results

As the next step in our analysis, we turn to the recorded EEG data. We analyzed the EEG signal first in the time domain, from -200 to 500 ms before and after gaze onset. As an example, we first focus on occipital electrodes to investigate visual evoked potentials (Luck, 2014). Using the classified gaze onsets, Figure 9A shows the fERP for one individual and averaged over all subjects. Overall, 3414 (IQR = 3050 - 4067) trials went into the calculation of single-subject ERPs. While minor between-subject variations were visible at the highest point for the P100, an average ERP across subjects revealed the shape of the P100, similar to literature (e.g., Luck, 2014). It was, however, visible that the underlying data and average fERPs are noisy, potentially caused by the movement and wearing of the head-mounted display during the experiment or by an unexact gaze classification. Investigating the distribution of the fERPs for all electrodes for one subject showed an expected pattern of high ERP, specifically P100, responses in the occipital electrodes (see Fig. 9B and 9C; Luck, 2014). Considering the time-sensitive nature of an ERP analysis, these results were a good indicator of the accuracy of determining gaze onsets. They indicate that combining eye-tracking data recorded in VR and EEG allows us to analyze the data in a meaningful way.

**Figure 9.**
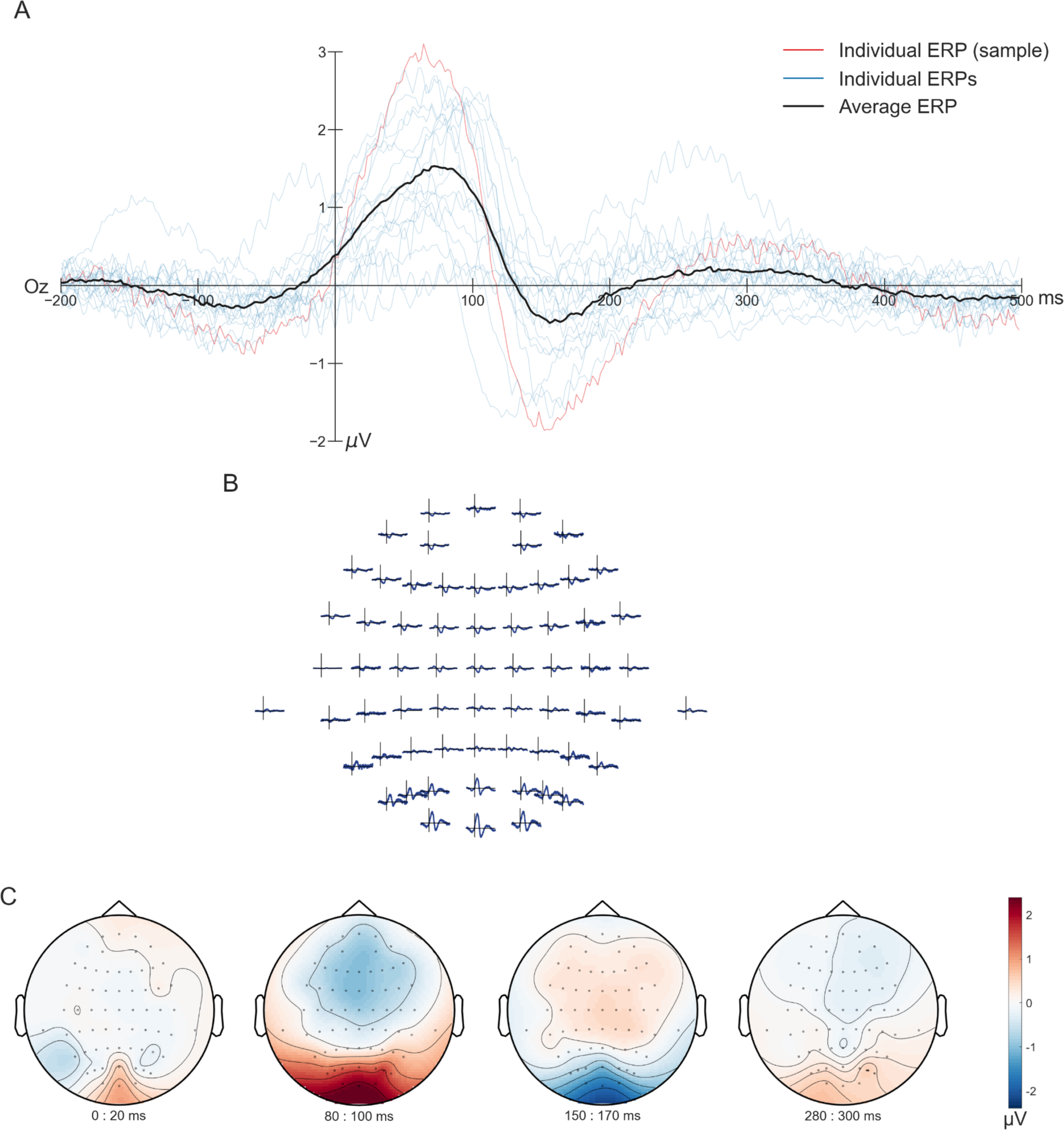
Fixation-Onset ERPs. **(A)** Here are fERPs at electrode Oz (64-channels; average reference) calculated for each subject (blue lines) and across subjects (black line), with the subject used as an example in other figures shown in red. The data was aligned to gaze onset and epoched from 200 ms before gaze onset until 500 ms after. **(B)** and **(C)** show the topographical distribution of the signal for one subject, the same as used for the single-subject eye-tracking plots. The time durations in **(B)** are identical to **(A)**, ranging from -200 to 500 ms. In **(C)**, the topographical distribution of four time points, 0-20 ms, 80-100 ms, 150-170 ms, and 280-300 ms, is presented. The topoplots (created using the default parameters of the FieldTrip ft_topoplotER (Oostenveld et al., 2011) function) for each time duration is an average across 20 ms.

As a next step, we analyzed fixation-onset ERSPs. For this purpose, spectrograms depicting the activation of one and all subjects at the electrode Oz using decibels (dB) as a unit were plotted (Fig. 10). The time was clipped from -500 ms to 800 ms, and the frequencies ranged between 2-45 Hz with 0.5 Hz increments. These spectrograms indicated an increased activation peaking at 100 ms followed by a decrease in power around 200 ms after fixation onset. The frequency range with a power increase was wider for the single subject than for all subjects, including theta, alpha, and beta bands. In contrast, the increase in mean activation across all subjects was only centralized to the theta and alpha bands. The decreased frequency range (5-35 Hz) did not differ between the single subject and all subjects. The fERSP results reveal a distinct pattern after gaze onset characterized by a power increase peaking 100 ms post-gaze onset, subsequently followed by a power decrease in the next 100 ms. These temporal power perturbations suggest that the experimental setup and recorded data are sufficiently accurate to yield significant results for high-quality analysis.

**Figure 10.**
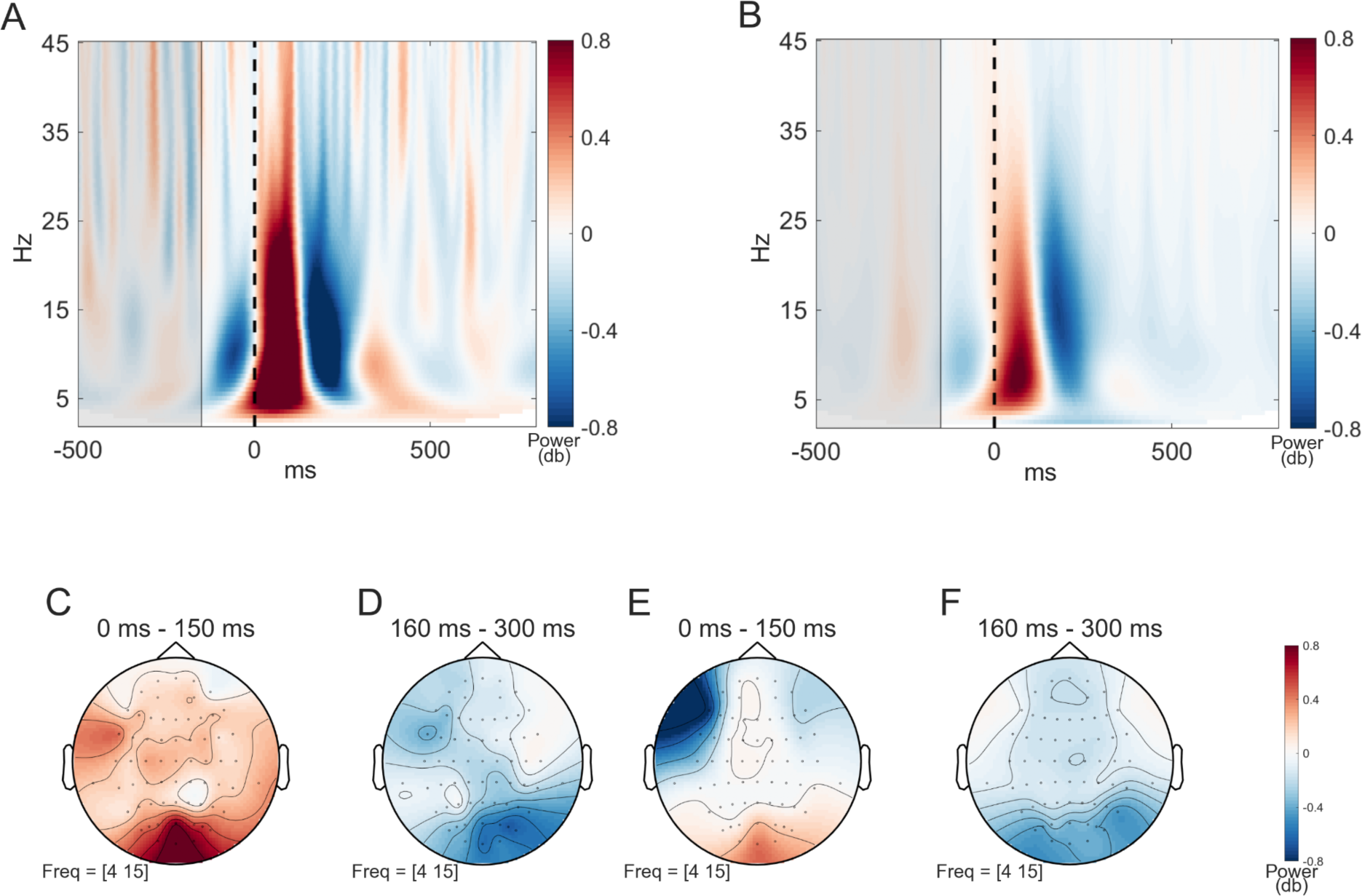
Power of fERSP (Fixation-Onset Event-Related Spectral Perturbations) **(A)** and **(B)** show the spectrograms of fERSPs at Oz (plotted using the FieldTrip ft_singleplotTFR function (Oostenveld et al., 2011); baseline = db). Time in seconds is given on the x-axis and the frequencies are on the y-axis. Warmer colors indicate an increase, and colder colors a decrease. The vertical dashed line marks the gaze onset at 0s. The baseline period ranged from -500 to -200 ms and is highlighted in gray. The data is averaged across all trials for one single subject at **(A)** and all subjects at **(B)**. In both spectrograms, an increase at around 100 ms followed by a decrease at approximately 200 ms can be observed. However, in a single subject **(A),** this increase’s frequency band is observed to be wider than the average of all subjects **(B)**. **(C)** and **(D)** show the scalp topographies for the frequencies 4-15 Hz for the single subject and **(E)** and **(F)** the corresponding topographies for all subjects. The plots **(C)** - **(F)** are created using the default parameters of the FieldTrip ft_topoplotTFR function (Oostenveld et al., 2011). The two different time intervals are chosen to depict the increase and the decrease in the power that would be comparable in both spectrograms - **(A)** and **(B)** - and they are 0-150 ms and 160-300 ms. **(C)** and **(E)** show both a power increase in the occipital electrodes for the first 150 ms after gaze onset, while for the second time interval, **(D)** and **(F)** show both a power decrease in the occipital lobe with a slight tendency to the right hemisphere.

A topographic analysis of the fERSP data was conducted to explore the spatial distribution of neural activity further. This analysis focused on the two-time windows with the strong power perturbations in the theta and alpha frequency bands (4-15 Hz) identified by the spectrograms for each condition. The first time interval extended from gaze onset to 150 ms post-gaze onset, while the second time interval covered 160-300 ms post-gaze onset. During the first time window, an increase in power was observed over the occipital electrodes, with a more pronounced power increase for the single subject (Fig. 10C) compared to all subjects (Fig. 10E). In the second time window, a consistent and substantial power decrease was observed over the occipital electrodes for a single subject and all subjects (Fig. 10D and 10F). Additionally, there was a slight tendency for the power changes to the right occipital hemisphere. The topographical maps suggest an occipital dominance of the power perturbations in the theta and alpha frequency bands during these time windows, aligning with previous literature (e.g., Cohen, 2014).

Finally, we assessed the time sensitivity and the potential influence of temporal uncertainty for both fERPs and fERSPs. To achieve this, we compared the Pearson correlation coefficients between fERPs and fERSPs, spanning the same time interval (-200 to 500ms), to individual trials with no shift, trials with a shift of minus one eye-tracking sample compared to gaze onset, and trials with a shift of plus one sample. Specifically, we compared the average fERP and fERSP with EEG trials using the same data as was used to calculate the average fERP/fERSP, once by setting the EEG trial onset to one eye-tracking sample in the past compared to the actual classified gaze onset, and once by taking an EEG trial onset one eye-tracking sample in the future compared to the actual gaze onset. With our eye-tracking sampling rate of 90 Hz, the two latter trial onsets were therefore shifted by 11 ms either into the future or past compared to the original trial onset. In the results of only one subject, we can see differences in the correlation coefficients between ERPs and ERSPs (Fig 11A). While the correlation coefficients and their distributions changed with the shift conditions for ERPs (Fig 11A & B), the correlation for ERSPs did not change as much. Yet, the median ERSP correlation coefficients were much higher than those of the ERPs (Fig 11A & C; the no-shift median for the ERP = 0.069 and for ERSP = 0.596). Across subjects, these observations are confirmed. The median correlation coefficients of the ERPs had a high between-subject variance and visible differences between all three shift conditions, with the no-shift having the highest correlation for all subjects. A repeated measure ANOVA and post hoc Bonferroni correction revealed a significant difference between the no-shift condition and both shifts for the ERPs (p-value no-shift - pos-shift = 0.000; p-value no-shift - neg-shift = 0.000) while no significant difference between the two shift conditions. Like ERPs, the median correlation coefficients of the ERSPs had a high between-subject variance, yet no difference between the shift conditions could be observed. A repeated measure ANOVA confirmed this, revealing no significant differences between the conditions. The assumption of sphericity for both ANOVAs was validated using the Mauchly test. Finally, the mean ERSP correlation for the no-shift condition across subjects was 0.596, close to .1 higher than the mean correlation coefficients of the ERPs. These results demonstrated that ERSPs are less time-sensitive than ERPs and have a smaller variance across trials, and are therefore better suited to study behavior in free-viewing experiments.

**Figure 11.**
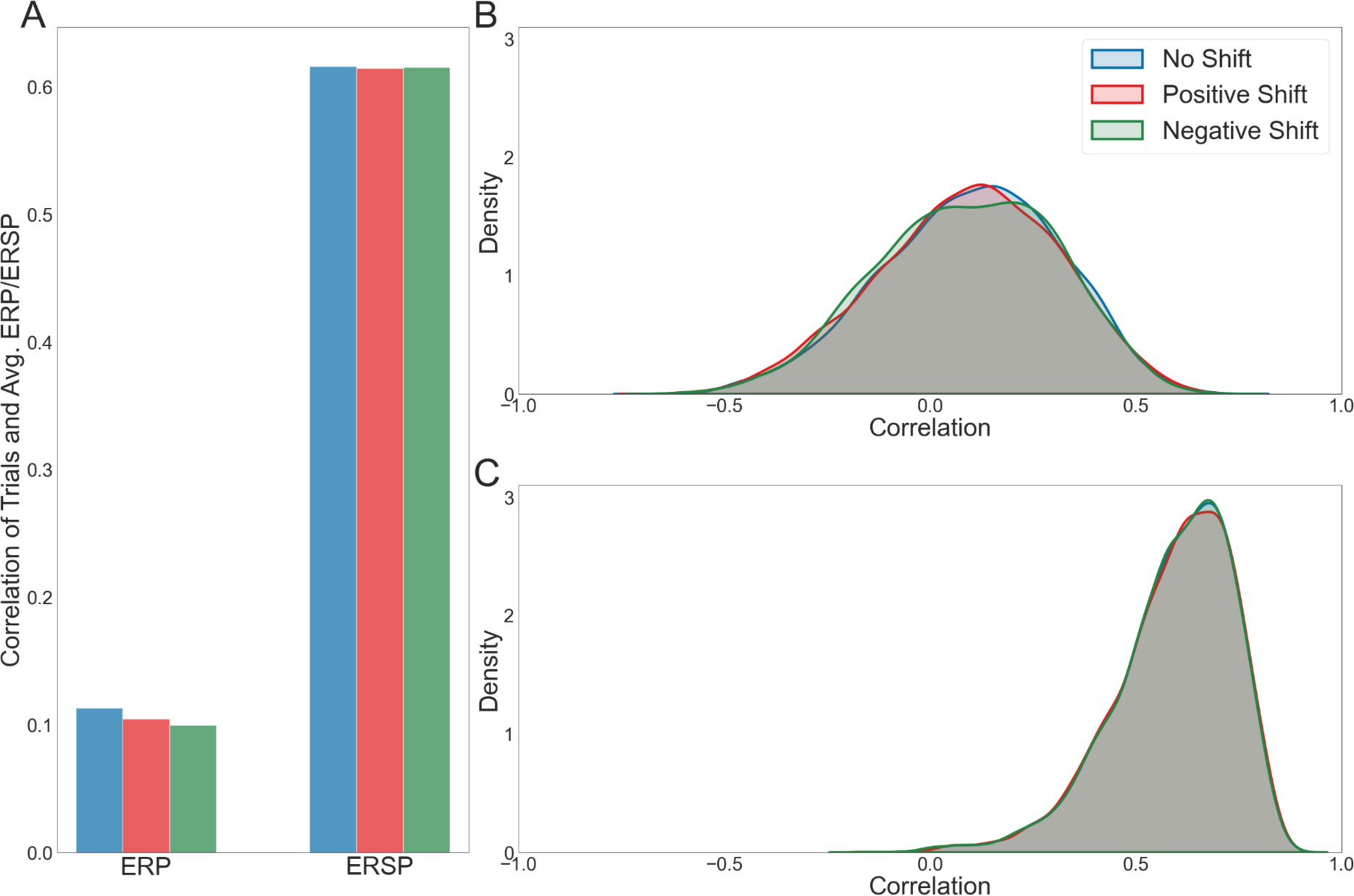
Correlation of Individual Trials with the Averaged fERP and fERSP. The Pearson correlation coefficients between each trial’s EEG signal and each subject’s average ERP and ERSP were calculated for channel Oz to compare the time sensitivity of the two measures. For this step, the data were high-pass filtered to 5 Hz as this analysis did not aim to investigate the effects of low-frequency drifts. The procedure was completed using trials with initial trial onsets classified with the eye-tracking algorithm (blue) and taking the trial onset as one eye-tracking sample before the originally classified gaze onset (red, in the middle) and one sample after the classified gaze onset (green). Both shifts were correlated with the average ERP and ERSP without a shift. In **(A)**, we provide the median correlation coefficients for all three conditions for one subject, the same as used in previous plots. The three left bars are the correlation coefficients for the ERPs and the right for the ERSPs. Correspondingly, **(B)** and **(C)** display the density distribution of the correlation coefficients for this subject, with **(B)** displaying the ERP results and **(C)** the ones for the ERSPs.

### Comparing the classification of our algorithm with hand-labeled data

As we do not have ground truth for our dataset, we assess the accuracy of the algorithm classification using hand-labeled data. At first, we compared the data on a sample-by-sample level, i.e., comparing how often a sample is classified by either procedure as being part of a gaze, saccade, invalid segment, or an outlier. The results can be seen in Figure 12A. We observe a high congruence of gazes. Interestingly, the data differs for the saccades, with roughly as many hand-labeled samples classified as saccades as gazes. For gazes and saccades, while the order of magnitude is the same between the hand-labeled data and the algorithm-classified data, the number of gazes and saccades between both differ. For the algorithm, we have a total of 4917 gazes and 4799 saccades; for the hand-labeled data, we have 6793 gazes and 6744 saccades. As the timing of the gaze onsets is relevant for the current purpose, we additionally assessed the temporal shift in gaze onsets of the algorithm compared to the hand-labeled data (see Fig 12B). Comparing gaze onsets up to +/- 110 ms revealed a median shift of 0.0 and an IQR = -1 - 3. It is important to note, however, that we had a total of 846, or 17.21%, algorithm-labeled gaze onsets that were further apart from hand-labeled gaze onsets than 110 ms. Besides the samples that could not be matched to a hand-labeled event, most gaze onsets were similar in time, as indicated by our median. However, the IQR demonstrates that our algorithm tends to define gaze events slightly later than the hand-labeled data.

**Figure 12.**
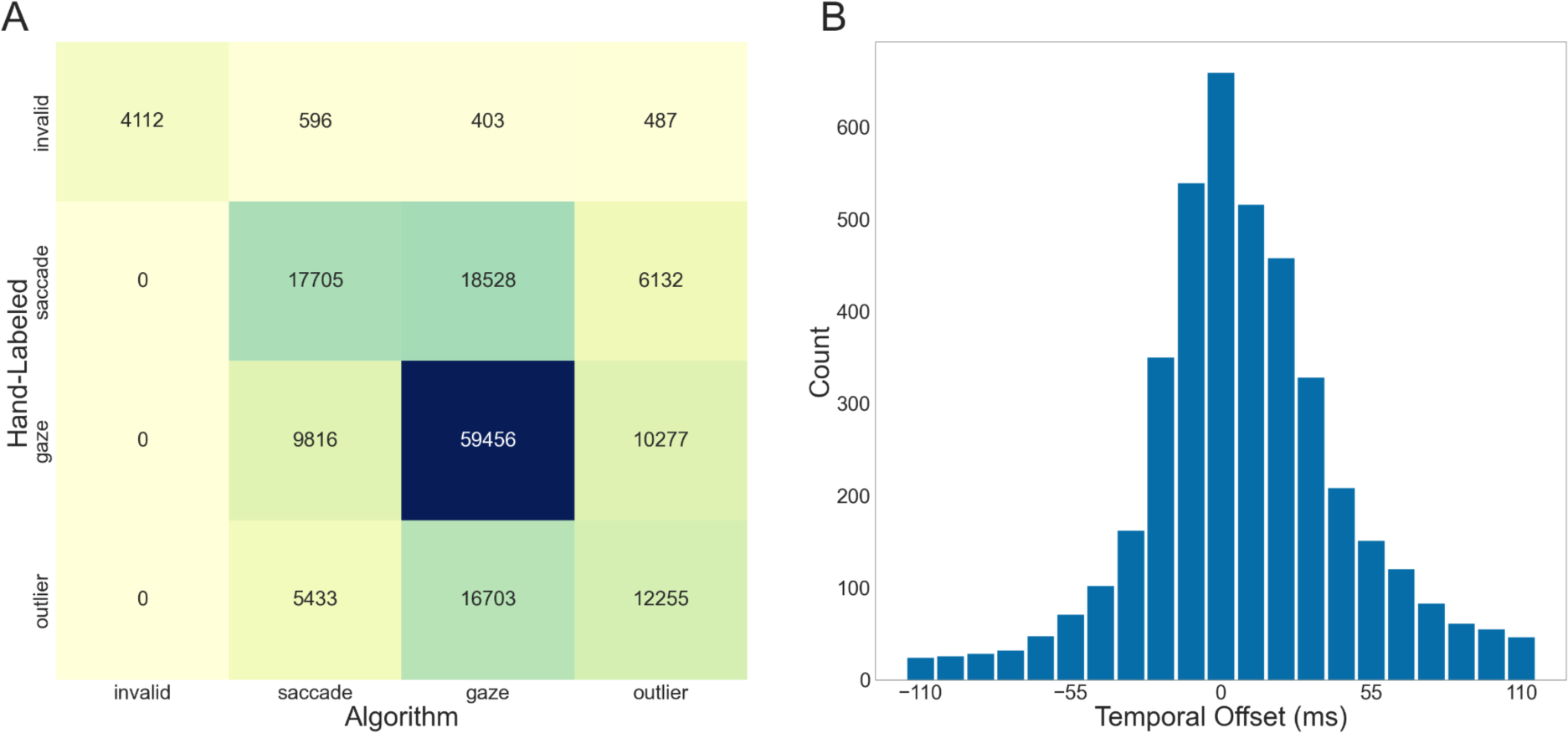
Comparison of the algorithm classification with hand-labeled data. **(A)** visualizes the sample-by-sample comparison of the labels assigned by the classification algorithm (x-axis) and by hand-labeling. In **(B)**, the temporal offset of the algorithm-defined gazes is compared to the hand-labeled data. The bar at 0 ms represents the amount of data where the algorithm-defined gaze onsets align with the hand-labeled onsets. The other bars correspond to the temporal shift in either direction. For example, the bar at -55 ms shows the number of gazes where the algorithm onset started 55 ms before the hand-labeled onset. If the gaze onset difference in the negative and positive direction was identical, we added ½ for both. As a note, the plot was created using frames, so the ‘temporal offset’ labels contain small variations in their timing.

## Discussion

The present study aimed to classify eye-movement data recorded in virtual reality to generate fixation-onset ERPs and ERSPs. Using our eye movement classification algorithm, we showed that we could accurately classify eye movements of three-dimensional free-exploration data and that we can generate fERPs and fERSPs, proving that combining EEG and free-viewing virtual reality setups is possible. We investigated the classification quality using our modified version of a velocity-based classification algorithm (Dar et al., 2021; Keshava et al., 2023; Voloh et al., 2020), correcting for subject movement in the virtual environment. Furthermore, we compared two data-segmentation methods dealing with varying noise levels across a long recording. The two methods exhibited a solid performance with only minor differences visible. As the different measures revealed no relevant significant differences, we consider the data-driven method (Dar et al., 2021) to elicit a fully adequate description of our data. Therefore, our classification algorithm’s final implementation is a combined and adapted version of the MAD saccade (Voloh et al., 2020) and REMoDNav algorithm (Dar et al., 2021), correcting for translational movement in a three-dimensional environment.

The recorded data used to classify gazes and saccades was clean, and both the gaze vectors and hit points accurately capture the eye movements. Our correction of translational movement, which improved the data quality, resulted in very clean eye-tracking data, comparable to results in the literature (Duchowski, 2017). Head movements and, to a smaller degree, eye movements predominantly occurred within the horizontal rather than the vertical direction; subjects rarely tilted their head upwards. The displayed behavior could have directly been caused by our instructions not to rotate the heads up and down. Another explanation for the observed behavior could be an environmental contribution: Most objects of interest were at participants’ eye height and might not have elicited further head or eye movements. Whether the results were caused by either one of these factors or a combination of both cannot be determined based on our data alone. However, similar results of higher variability in the horizontal compared to the vertical eye-in-head direction could be observed in real-world free-viewing studies (Foulsham et al., 2011; ’t Hart & Einhäuser, 2012), and it has been shown that subjects’ heading direction was aligned with selective visual attention (Thom et al., 2023). Overall, the three-dimensional eye-tracking data recorded using virtual reality was comparable to literature findings and suitable for classifying separate events.

After classifying our eye-tracking data into gazes and saccades, several measures can be employed to investigate the quality of the classification. As a first measure, the average duration and duration distributions of saccades and gazes are comparable to the literature (Dar et al., 2021; Llanes-Jurado et al., 2020; Nyström & Holmqvist, 2010). We have skewed, non-Gaussian distributions for both gazes and saccades. Our saccade durations are slightly longer than previous data presented in the literature (Dar et al., 2021; Nyström & Holmqvist, 2010), most likely due to our lower sampling rates. In order to exclude potential outliers, saccades with a duration of only a single sample were rejected. The finite sampling rate resulted in a relatively high observable minimum saccade duration and, therefore, a longer median and mean duration. The mean gaze duration, on the other hand, aligned with previously reported data (e.g., Duchowski, 2017; Llanes-Jurado et al., 2020). It is comparable between subjects, proving that the event classification mostly works as intended. Interestingly, this study’s between-subject differences are smaller than those of other studies (e.g., Nyström & Holmqvist, 2010). It is hard to differentiate if this results from our free viewing paradigm or is based on the three-dimensional data we collected with the relatively lower sampling rate. Similarly, the velocity distributions across eye-movement event sequences and subjects follow a pattern expected when using a velocity-based algorithm: We see high velocities before and comparable lower velocities after gaze onset (Nyström & Holmqvist, 2010). The wide distribution of saccade peak velocities with the highest density close to gaze onset mimics the distribution of saccade durations. It, therefore, provides additional support for the known underlying saccade velocity profile (Harris & Wolpert, 2006). As velocity and dispersion distributions in relation to gaze onset are often related (Duchowski, 2017), our dispersion distributions supply additional support for expected underlying saccade and gaze profiles. In contrast, while the overall performance of the classification algorithm is similar to the hand-labeled data, the comparison raises two main points of consideration. First, the two methods resulted in a different number of events, with many hand-labeled saccades being classified as gazes. This difference might have been caused by the comparatively long (Dar et al., 2021) minimum saccade and gaze durations, an effect of the low sampling rate. Additionally, investigating the event-onset shifts revealed a tendency of algorithm-based event onsets to start after hand-labeled gaze onsets. This discrepancy could be caused by the different classifications of post-saccadic oscillations (Dar et al., 2021). While our algorithm might classify them as part of a saccade, the hand-labeled data might already consider them part of the gaze. Therefore, while our classified events follow many patterns observed in the literature, demonstrating that our classification algorithm might work as intended, it also highlights room for improvement.

Trusting in the current classification, gaze onsets were utilized for fERPs and fERSPs to investigate the combination of EEG and virtual reality recordings. The time-sensitive aspect of the EEG signal can be used to easily examine the signal quality (Cohen, 2014; Luck, 2014) and, consequently, the quality of the definition of saccade offsets and gaze onsets. Investigating fERPs or fERSPs based on data recorded with virtual reality poses the challenge of dealing with the low eye-tracking sampling rates (Duchowski, 2017). According to Luck (2014), temporal inaccuracies of ± 10ms are acceptable, indicating that a stable frame rate of 90 Hz is most likely suitable for this analysis. Our data confirms this observation, as the generated fERPs show a similar time course to that reported in previous literature (Luck, 2014). As an important note, we did not yet control and correct for temporally overlapping events (Ehinger & Dimigen, 2019; Gert et al., 2022; Henderson et al., 2013) which might affect the overall shape of our fERPs. Specifically, for shorter fixation durations, the following fixation can occur earlier compared to longer fixations, and as a result, the early neuronal components of the following fixation will be superimposed on the late components of the preceding fixation (Dimigen et al., 2011; Henderson et al., 2013). Furthermore, a possible modulation of the overall shape of the fERPs due to the relatively high high-pass filter (Tanner et al., 2015) cannot be disregarded. This consideration might be particularly important when investigating modulations of ERPs caused by different experimental conditions (Tanner et al., 2015). Besides the fERP analysis, using a Morlet wavelet transformation, the time-frequency analysis yielded promising results with changes in the activity at 100 ms and 200 ms after the stimulus onset, similar to the fERP components. Our results from the time-frequency analysis yielded activity in the theta to beta band frequencies but only very low activity at the gamma frequencies. One reason for the low gamma-band activity could be the saccadic spike artifacts, which affect experimental setups with non-stationary stimulation (Hipp & Siegel, 2013), such as in our study. However, our study differs from this publication in two aspects. First, we looked at the activity at the occipital rather than the parietal electrodes (Hipp & Siegel, 2013). Second, to be comparable with the fERP analysis, our time-frequency analysis favored a better time resolution and, therefore, lacked precision in the frequency resolution. As a result, to investigate the gamma frequencies further, the analysis should be optimized for that objective. Importantly, differences in the correlation between the activity at trial onsets without and trial onsets with temporal jitter only influenced fERPs and not fERSPs, meaning fERSPs are less affected by temporal uncertainty, confirming findings from the literature (Cohen, 2014). This indicated that in recordings challenged with temporal inaccuracies or lower sampling rates, fERSPs might be the better-suited analysis. Besides temporal uncertainty, the correlation coefficients were higher for fERSPs than fERPs, indicating a lower between-trial variance and, thus, fewer trials required to estimate a sound time-frequency analysis. For experiments using a free-viewing and free-exploration context, this reduces the burden of precisely reproducing experimental conditions with a large number of trials and might, therefore, be the preferred method. Combining eye-tracking data recorded in virtual reality and classified using our algorithm has a high enough temporal precision to generate fERPs and fERSPs, yet for free-viewing experiments, fERSPs might provide more accurate results.

Combining EEG and virtual reality in a free-viewing setup raises the question of possible influences on the signal quality, especially since, even after careful preparation, our data was quite noisy for many subjects, some of which could not be included in the final data analysis. The influence of head movements and HMDs on the quality of EEG signals has been investigated in various studies, including our own experiences (Izdebski, et al., 2016; Oliveira et al., 2016; Wang et al., 2020; Weber et al., 2021). It is known that vertical and, to a smaller degree, horizontal head movements greatly affect the signal-to-noise ratio (Luck, 2014; Oliveira et al., 2016; Tauscher et al., 2019), yet preprocessing steps, such as ICA can handle these motion artifacts (Oliveira et al., 2016). Furthermore, studies have demonstrated that it is possible to collect EEG data while subjects are wearing an HMD without reducing the signal quality (Izdebski et al., 2016; Wang et al., 2020; Weber et al., 2021). It is clear that specific care has to be taken, but in our own work, we did not observe a negative impact of combining these two methods (Izdebski et al., 2016; Oliveira et al., 2016).

Finally, when combining EEG and VR in free-viewing and exploration studies, some general points need to be considered. Integrating different methodologies while subjects move freely can lead to high data drop-out. In the current study, almost half of all subjects could not be included in the final data analysis. Other than noisy EEG signals, motion sickness was an issue for many subjects. Motion sickness can be dealt with through different means, e.g., through short recordings, active movement in the virtual and simultaneously in the real world (Clay et al., 2019), and with the help of motion sickness tests at the beginning of the recording. Overall, planning to record more subjects than needed is essential. Another point to consider is the accurate alignment of timestreams. Combining eye-tracking and EEG data in long recordings without predefined trials requires a constant frame rate, which can challenge specific virtual reality setups (e.g., Walter et al., 2022). Finally, velocity-based algorithms generally require high sampling rates (Duchowski, 2017; Larsson et al., 2013), so should the sampling rate fall below 90 Hz, another quality check of the proposed algorithm and a comparison between the two data segmentation methods might be needed. Altogether, while it is possible to combine EEG and eye-tracking data recorded in a free-exploration, virtual reality experiment, specific issues can arise that make the combination of both challenging.

In this paper, we implemented an algorithm that is suitable for determining gaze onset events. During the early phases of the project, we pursued the preferable choice of using already existing tools. There are a number of tools available to classify eye movements for static 2D images (e.g., Dar et al., 2021); however, we found that dynamic scenes created by the 3D environment, as well as allowing subjects to move, caused a problem. So, step by step, we modified and appended existing algorithms to this more demanding scenario (Dar et al., 2021; Keshava et al., 2023; Voloh et al., 2020). Specifically, adding EEG into the mix considerably increased the demand for precision (Luck, 2014). With the current algorithm, we have a workable solution, yet it does not mean that this is the end of the development. Specifically, differences between the hand-labeled data and the algorithm-defined classifications raise points of improvement, such as the different number of events or the timing of event onsets. Delving further into these differences might refine the classification. Additionally, further improvements on the algorithm to allow a subsample precision in defining fixation onset would be desirable. Such a solution would decouple the EEG signal sampled at a high frequency from measuring the eye movements typically sampled at a much lower frequency in a VR setup. Overall, however, we consider the current version to perform reasonably well enough for the current purpose of combining EEG and eye-tracking data.

In conclusion, this project aimed to investigate the possibility of combining eye-tracking data recorded in virtual reality with EEG data in a free viewing study. We modified and tested a version of the MAD saccade (Keshava et al., 2023; Voloh et al., 2020) and REMoDNav (Dar et al., 2021) algorithm while additionally correcting for subjects’ translational movement within the virtual scene. The behavioral results indicate an accurate classification of eye movements into gazes and saccades. Using the gaze onsets as trial onsets, we generated fixation-onset ERPs and ERSPs, with fERSPs being less time-sensitive and requiring fewer trials to be estimated; they are therefore better suited for more naturalistic experiments. These results provide a methodological basis for combining different techniques, such as EEG and eye-tracking, in more realistic free-viewing and free-exploration virtual reality studies. Therefore, as virtual reality allows for investigating phenomena in more naturalistic settings without losing the replicability and control usually provided by traditional laboratory studies, this opens the possibility for many questions to be answered in the future.

### Open Practices Statement

The data and the code for the experiment reported here are unavailable but will be made publicly available on the Open Science Framework before publication. The experiment was not preregistered.

